# Satb2 regulates proliferation and nuclear integrity of pre-osteoblasts

**DOI:** 10.1101/574863

**Authors:** Todd Dowrey, Evelyn E. Schwager, Julieann Duong, Fjodor Merkuri, Yuri A. Zarate, Jennifer L. Fish

## Abstract

Special AT-rich sequence binding protein 2 (Satb2) is a matrix attachment region (MAR) binding protein. Satb2 impacts skeletal development by regulating gene transcription required for osteogenic differentiation. Although its role as a high-order transcription factor is well supported, other roles for Satb2 in skeletal development remain unclear. In particular, the impact of dosage sensitivity (heterozygous mutations) and variance on phenotypic severity is still not well understood. To further investigate molecular and cellular mechanisms of Satb2-mediated skeletal defects, we used the CRISPR/Cas9 system to generate *Satb2* mutations in MC3T3-E1 cells. Our data suggest that, in addition to its role in differentiation, Satb2 regulates progenitor proliferation. We also find that mutations in *Satb2* cause chromatin defects including nuclear blebbing and donut-shaped nuclei. These defects may contribute to a slight increase in apoptosis in mutant cells, but apoptosis is insufficient to explain the proliferation defects. *Satb2* expression exhibits population-level variation and is mostly highly expressed from late G1 to late G2. Based on these data, we hypothesize that Satb2 may regulate proliferation through two separate mechanisms. First, Satb2 may regulate the expression of genes necessary for cell cycle progression in pre-osteoblasts. Second, similar to other MAR-binding proteins, Satb2 may participate in DNA replication. Deficiencies in either of these processes could reduce the pace of cell cycle progression and contribute to nuclear damage. We also hypothesize that Satb2-mediated proliferation defects may be buffered in some genetic backgrounds, which provides some explanation for differences in severity of skeletal defects. Further elucidation of the role of Satb2 in proliferation has potential impacts on our understanding of both skeletal defects and cancer.

## 1. Introduction

Bone formation in development and regeneration requires proliferative expansion of pre-osteoblasts and their subsequent differentiation. These two processes rely on the function and temporal coordination of multiple transcriptional regulators. In particular, *Runx2* and *Sp7/Osx* are essential to osteogenesis as deletion of either impairs osteoblast differentiation and eliminates bone formation (Komori *et al.*, 1997; Nakashima *et al.*, 2002). *Runx2* regulates the expression of *Sp7/Osx* as well as the bone matrix proteins *Ocn*, *Bsp*, and *Opn* (Komori, 2005). *Runx2* also promotes the expression of *Atf4*, a regulator of terminal differentiation in osteoblasts (Yang *et al.*, 2004).

The function of these transcription factors is augmented by Satb2 (Special AT-rich sequence binding protein 2), a protein that binds to nuclear matrix-attachment regions (MARs). It is thought that Satb2 organizes chromatin looping at MARs such that the expression of cell-type specific differentiation genes may be enhanced (Britanova *et al.*, 2005). In mice, loss of Satb2 results in skeletal hypoplasia and reduced expression of osteogenic genes including *Ocn* and *Bsp* (Dobreva *et al.*, 2006). In contrast, *Satb2* over-expression was found to promote bone formation and increase *Ocn* and *Bsp* expression (Gong *et al.*, 2014; Zhang *et al.*, 2011). These results are consistent with skeletal malformations in humans with *SATB2*-associated syndrome (SAS), which include bowing tibias, low bone mineral density with high fracture rate, osteoporosis, cleft palate, micrognathia, and maxillary hypoplasia (Boone *et al.*, 2016; Leoyklang *et al.*, 2007; Zarate and Fish, 2017; Zarate *et al.*, 2018a; Zarate *et al.*, 2018b).

One role of Runx2 in pre-osteoblasts is to down-regulate the miR cluster 23a/27a/24-2. These miRs target the 3’ UTR of *Satb2*, and thus Runx2 alleviates the repression of *Satb2* (Hassan *et al.*, 2010). These and other data have placed Satb2 downstream of Runx2 in the osteogenic differentiation program, with mutations in *Satb2* largely thought to impair the expression key genes involved in osteoblast maturation (Dobreva *et al.*, 2006; Gong *et al.*, 2014; Zhang *et al.*, 2011). Although the majority of investigations into Satb2 function in osteogenesis have focused on its role in gene regulation, loss of Satb2 has also been reported to cause apoptosis in neural crest progenitors of the jaw (Britanova *et al.*, 2006; Fish *et al.*, 2011).

To further explore molecular and cellular mechanisms underlying *Satb2*-mediated defects in osteogenesis, we used the CRISPR/Cas9 system to generate mutations in the *Satb2* locus in MC3T3-E1 cells, which are clonal pre-osteoblasts that have been widely used as a model of osteoblast differentiation (Xiao *et al.*, 1997). Several modified colonies with different combinations of *Satb2* alleles producing a range of *Satb2* expression levels were selected for experimental analysis. Consistent with previous reports, we observe altered osteogenic gene expression in *Satb2* mutants. However, we also find that reduction in *Satb2* expression, even that of true heterozygotes, results in reduced pre-osteoblast proliferation. Further, mutations in *Satb2* cause nuclear aberrations including nuclear blebbing and donut-shaped nuclei.

Our data point to a novel role for Satb2 in mediating pre-osteoblast proliferation, indicating that Satb2 is involved in both proliferative expansion of pre-osteoblasts and their subsequent differentiation. Previous studies failing to observe proliferation defects in *Satb2* mutants may reflect buffering of this process in some genetic backgrounds. The potential role of genetic background in regulating susceptibility to Satb2-mediated proliferation defects may help explain variation in penetrance and severity of skeletal defects in SAS patients. Participation in both proliferation and differentiation processes makes Satb2 an interesting target for bone regeneration and osteoporosis therapies (Guo *et al.*, 2016; Zhang *et al.*, 2011). Finally, our observation that mutations in *Satb2* both reduce proliferation and cause nuclear aberrations has implications for understanding its dichotomous potential for both tumor suppression and oncogenesis (Chen and Costa, 2018).

## 2. Materials and Methods

### Cell lines

Mouse osteoblast precursor cells (MC3T3-E1 Subclone 4) obtained from ATCC were grown in αMEM supplemented with 10% FBS and Penicillin/Streptomycin. Passage numbers higher than 20 were discarded.

### Generation of mutant lines using CRISPR/Cas9

To generate indel mutations near the start codon of *Satb2* (Fig. 1A), we used a previously validated guide RNA, *Satb2*-272, ATCGGAGCAGCATGGAGCGG (Shinmyo *et al.*, 2016). The guide RNA was cloned into pSpCas9(BB)-2A-Puro (PX459) V2.0 (Addgene plasmid # 62988), as previously described (Ran *et al.*, 2013). Cells were transfected using the NEON transformation system (Invitrogen) using one 1200V pulse and a 30 ms pulse width and otherwise following the manufacturer’s protocol. 48-hours after electroporation, 3 μg/mL Puromycin was added to the cells for 24 hours. Surviving cells were diluted to 2.5 cells/ml and seeded into 96-well plates at 0.5 cells per well to obtain single cell colonies. Single colonies were grown to confluency and then expanded.

**Figure 1:**
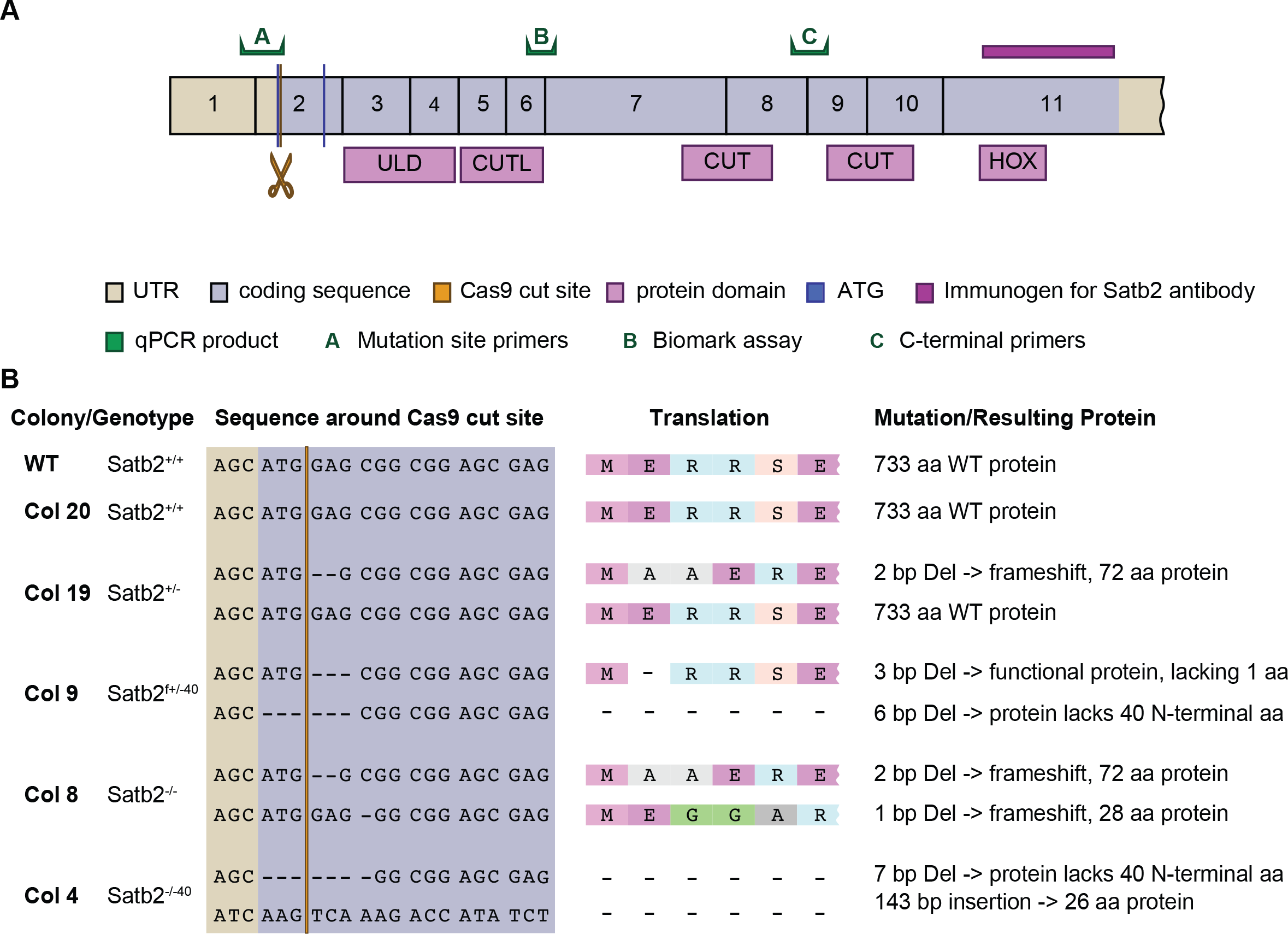
Overview of CRISPR strategy and outcomes. A) Diagram of Satb2 protein with the location of the Cas9 cut site, qPCR primer sets, and the binding region of the antibody used in this study highlighted. B) Colony cell lines used in this study are described. The colony number, *Satb2* genotype, cut site DNA sequence and translation, as well as protein product produced are listed.

### Analysis of mutant cell lines

DNA was extracted from single cell colonies using the ZR Duet DNA/RNA Miniprep Kit (Zymo). A 708 bp PCR product around the editing site of *Satb2-*272 was amplified using primers Satb2-seq-F ACTGGCCTGATCGTCTATCA and Satb2-seq-R GCCAGATCCTAGGTCTCTGT. PCR fragments were cloned into TOPO pCR4 (Invitrogen) and sequenced with M13 F+R primers using Sanger sequencing to determine the exact sequence of each allele.

### Differentiation assay

An overview of the differentiation assay is shown in Figure 6A. Cells were plated in a 12-well plate at 5 × 10^4^ cells per well. After 3 days, differentiation was induced (=T0) with Dulbecco’s modified Eagle Medium supplemented with 10% FBS, Penicillin/Streptomycin, 50 µg/ml beta-glycerophosphate, and 200 µM ascorbic acid. The media was changed every 2-3 days over the differentiation period (0-28 days). Mineralization was detected with 1% (w/v) Alizarin S pH 4.2 for 1 h and imaged on an Epson Perfection 4990 Photo scanner and an Olympus BX41 microscope.

### Immunohistochemistry

Cells were fixed in 4%PFA/PBS for 15 minutes, permeabilized with 0.1% Triton-X-100/PBS and then blocked in 5% FBS supplemented with 0.1% BSA for 1 h. For BrdU, cells were treated with an additional incubation in 2N HCl for 1 h at room temperature after the permeabilization. Primary antibodies were incubated overnight at 4°C in blocking solution at the following concentrations: mouse anti-Satb2 (SATBA4B10, SantaCruz) 1:300; mouse anti-alpha Tubulin (DM1A) 1:500; mouse anti-Lamin B1 (B-10, SantaCruz) 1:100; rabbit anti-Lamin A/C (Cell Signaling Technology) 1:100; rabbit anti-BrdU (Abcam) 1:50. Cells were imaged on a Nikon AR-1, a Leica Sp8, or a Zeiss Axiovert 200M microscope. Images were processed using the Fiji distribution of ImageJ and Adobe Photoshop CC 2018 (Schindelin *et al.*, 2012).

### Imaging flow cytometry

Cells for the flow analysis were plated at 4 × 10^5^ four days before the experiment. After trypsinizing, cells were resuspended in media and counted using a hemocytometer. Cells were spun down, resuspended in PBS to 1 × 10^6^, and incubated for 15 min at room temperature in the dark in 1x Annexin Binding Buffer (Invitrogen) supplemented with 2 µl Annexin V, Alexa Flour 488 conjugate (Invitrogen) and 0.5 µg/ml propidium iodide (Alfa Aesar). After a final centrifugation step, cells were resuspended in 100 µl 1x Annexin Binding Buffer and then analyzed immediately on an Amnis imaging flow cytometer with a 20x objective. Both Annexin V A488 and PI were excited using the 488 laser. Brightfield (channel 01 and 09, 435-480 nm and 570-595 respectively), Annexin V A488 (channel 02, 480-560 nm) and PI (channel 04, 595-642 nm) were measured and 10000 single cell events (see gating below) were counted and all events collected. Color compensation was achieved using single-stained samples that contained cells in which apoptosis was induced by incubating cells for 24-48h with 5 µM camptothecin. Focused cells were gated using the gradient root mean square of the bright field image, and bright field area and aspect ratio were used to gate single cells. Using fluorescence intensity of channel 02 (Annexin V A488) and channel 04 (PI), cells were divided into double negative (live) cells, Annexin V-positive cells (early apoptotic) and double positive cells (late apoptotic and necroptotic).

### qPCR

RNA was collected with the illustra triplePrep kit (GE healthcare) or TRIzol (Invitrogen). RNA was DNase-treated with the TURBO DNA-Free kit (Invitrogen) and quantified using a Qubit 3 Fluorometer (Invitrogen). cDNA was generated with 400ng of RNA with iScript RT Supermix for RT-qPCR (BioRad). GAPDH and Satb2 primers were from (Gong *et al.*, 2016), Satb2 cut (F: CCCCAGCCAGCCAAGTTTCA, R: GCCGGAGTCTGTTCACTACC), Runx-2 (F: ATTCGTCAACCATGGCCCAG, R: GAAACCCAGTTATGACTGCCC), Bsp/Ibsp (F: ATGGAGACGGCGATAGTTCC, R: ACACCCGAGAGTGTGGAAAG), Ocn/Bglap (F: TTCTGCTCACTCTGCTGACC, R: GCCGGAGTCTGTTCACTACC), Osx/Sp7 (F: GCCGGAGTCTGTTCACTACC, R: GCCGGAGTCTGTTCACTACC). Fold change was calculated using the delta-delta C(t) method (Livak and Schmittgen, 2001).

### Single Cell qPCR

Cells were trypsinized and at T14 or later also treated with collagenase type 1 (Worthington) for 15-30 min at 37°C. Cells were loaded onto a C1 Single-Cell Preamp integrated fluidic circuit (IFC) with 17–25 μm capturing well size. Primers for pre-amplification were obtained from Fluidigm (Delta Gene Assays, see Supplementary Table 1). After single-cell capturing, the inlets of the IFC were microscopically checked for the presence of single cells. Empty inlets or inlets with two or more cells were recorded and excluded from the qPCR analysis. Lysis, reverse transcription and preamplification was performed according to protocol. Single cell qPCR was performed using the Fluidigm BioMark system on 96.96 Dynamic Array IFCs for Gene Expression using Delta Gene Assays and Sso Fast EvaGreen Supermix with Low ROX (Bio-Rad). Single cell qPCR was performed using the Fluidigm BioMark system on 96.96 Dynamic Array IFCs for Gene Expression using Delta Gene Assays and 2X SsoFast EvaGreen Supermix with low ROX (Bio-Rad). For data-processing, the Fluidigm Real-Time PCR Analysis Software v4.5.2 was used to eliminate failed assays. The default quality threshold cutoff (0.65) was used to identify potential artifacts. The baseline correction was to linear, and the Ct threshold detection method was set to Auto (Detectors). The Fluidigm Singular Analysis Toolset v3.6.2 was used in *R* to analyze single cell gene expression. The statistical computing software *R* was used to identify patterns in gene expression.

## 3 Results

### 3.1 Satb2 mutant lines

We selected several CRISPR/Cas9-modified cell lines with different types of mutations for analysis. Our selected colonies include two heterozygous lines and two homozygous mutant lines (Fig. 1). Of the heterozygous lines, one is a true heterozygote with one WT allele and one loss of function allele (colony 19), while the other heterozygote has an allele missing a single amino acid at the N-terminal end (with predicted WT function) and an allele with a N-terminal 6bp deletion that includes the ATG (colony 9). Of the two homozygous mutant lines, one is a complete loss of function mutant with two alleles carrying frameshift deletions (colony 8). The other has one allele with a 7bp deletion and one allele with a 143bp insertion, both removing the ATG (colony 4).

*Satb2* mRNA and protein levels were evaluated in each of these lines by qPCR and Western blot, respectively. Using two sets of primers (A and C in Fig.1A), we find that no full-length mRNA is produced for any allele containing a mutation in the start codon (**SupFig. 1A**). Using primers directed against the C-terminal end of the gene, we find *Satb2* mRNA is produced in all our cell lines. However, some of this mRNA (especially that of colony 8) is predicted to form non-functional (frame-shifted) protein or not be translated at all. Western blot confirmed colony 8 is a loss of function mutant (**SupFig. 1B**). Similarly, colony 19 produces roughly half the amount of Satb2 protein relative to WT, as expected for a true heterozygote. Interestingly, we found that the short N-terminal deletions found in colonies 9 and 4 produce a protein. Although the start codon is deleted in both affected alleles, another ATG exists 120bp downstream. Our data are consistent with this allele producing a Satb2 protein missing the first 40 amino acids. These 40 amino acids are not part of any previously described functional domains (Fig. 1A).

Immunostaining using an antibody targeted against the C-terminal end of Satb2 provides results consistent with our Western blot analyses (Fig. 2). Importantly, the immunostaining data confirm the production of protein lacking the N-terminal 40 amino acids (**see C4 in** Fig. 2). Also of note is that in colony 9, which produces 2 Satb2 proteins of different sizes, the shorter protein appears to be preferentially produced (**SupFig. 1B**). This preference for producing the shorter protein has been observed in 4 separate Western blots.

**Figure 2:**
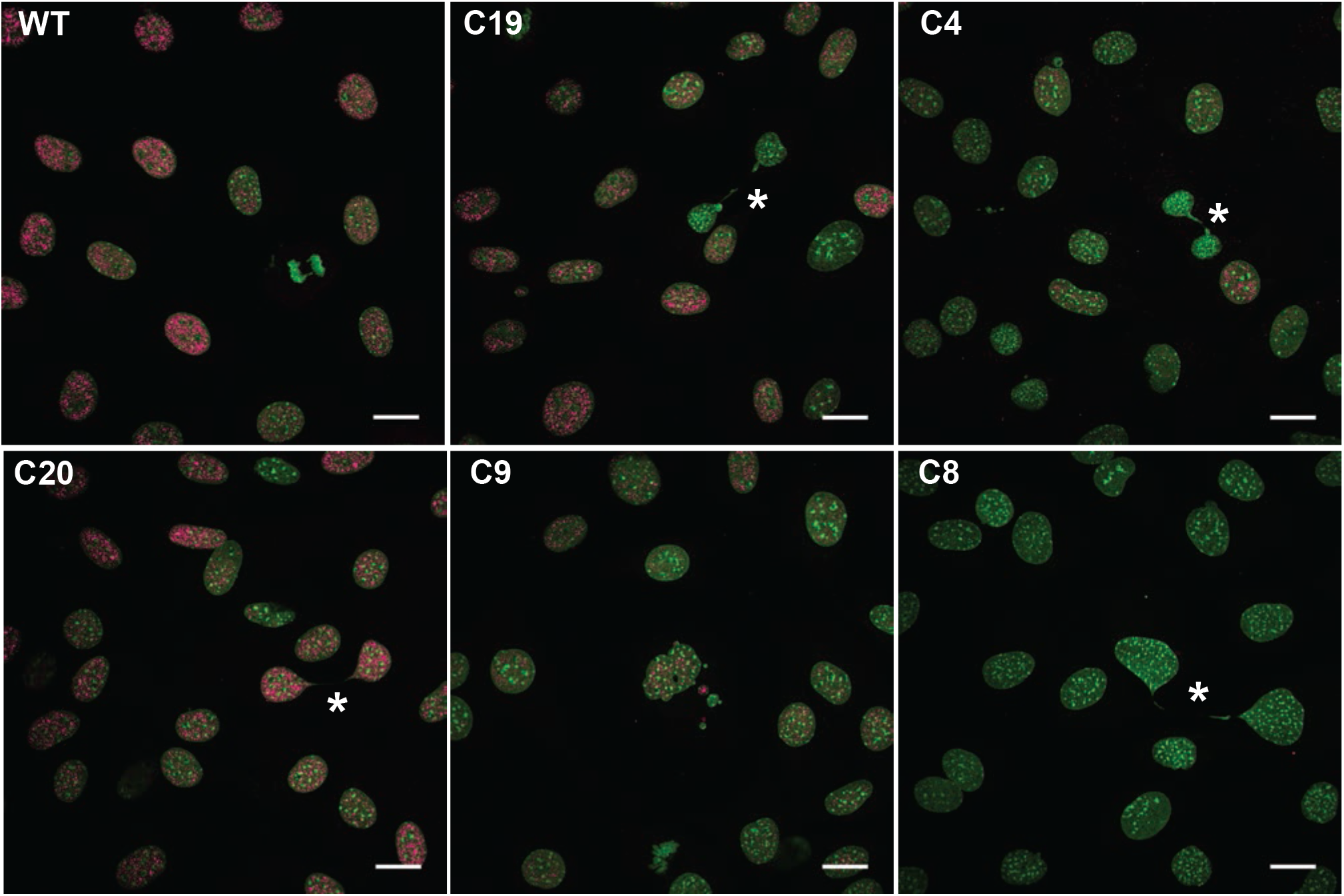
Mutations in Satb2 reduce protein levels. Confocal maximum intensity projections showing representative *Satb2* protein levels in undifferentiated (T0) cells of colonies used in this study. Note the presence of Satb2 protein in C4, indicating the production of a mutant protein, as well as the absence of protein in C8. DNA in green; Satb2 in pink. Asterisks highlight chromatin bridges. All scale bars represent 10 µm.

### 3.2 Satb2 and the cell cycle

When assessing Satb2 levels, we noticed that in all cell lines, Satb2 exhibits population variation (Fig. 2). In interphase cells, Satb2 ranges from intense to weak, although always associated with euchromatin. Satb2 is present in early mitosis, however it is no longer associated with chromatin, and is absent by late mitosis (Fig. 2A **and** 3A). As part of our experiments to investigate variation in osteogenic gene expression (described below), we performed single cell gene expression analyses. Interestingly, we found that in WT cells, *Satb2* expression levels cluster among cell cycle regulators rather than with genes associated with osteogenic differentiation (**SupFig. 2**). Notably, the relative expression of a sub-set of cell cycle genes associated with late G1 and S phases separated the cells into two main clusters (**SupFig. 2**). In particular, the genes *Cdc25a* (G1-S), *Cdc6* (S), *Cdk2* (G1-S), and *Cdk1* (S-M), along with *Satb2*, were highly expressed in one cluster relative to the other (**see rectangle, SupFig. 2**). In contrast, expression of genes associated with G1 such as *Cdcn1*, *Cdcn2*, *Cdk4*, and *Cdk6*, was more widely distributed among cells. These data suggest that *Satb2* is cell cycle regulated and/or regulates cell cycle genes.

To further investigate Satb2 variation during the cell cycle, we evaluated Satb2 levels in cells that had been treated with BrdU for 2 hours. We binned cells into 3 classes of Satb2 levels. Low to no Satb2 protein is observed in cells in anaphase through early G1 (see Fig. 2 **and** 3). BrdU positive cells, indicating cells in S and G2 have high levels of Satb2, however, high Satb2 levels are also observed in BrdU negative cells (Fig. 3). Taken together, these data suggest that Satb2 is most active from late G1 to late G2.

**Fig. 3:**
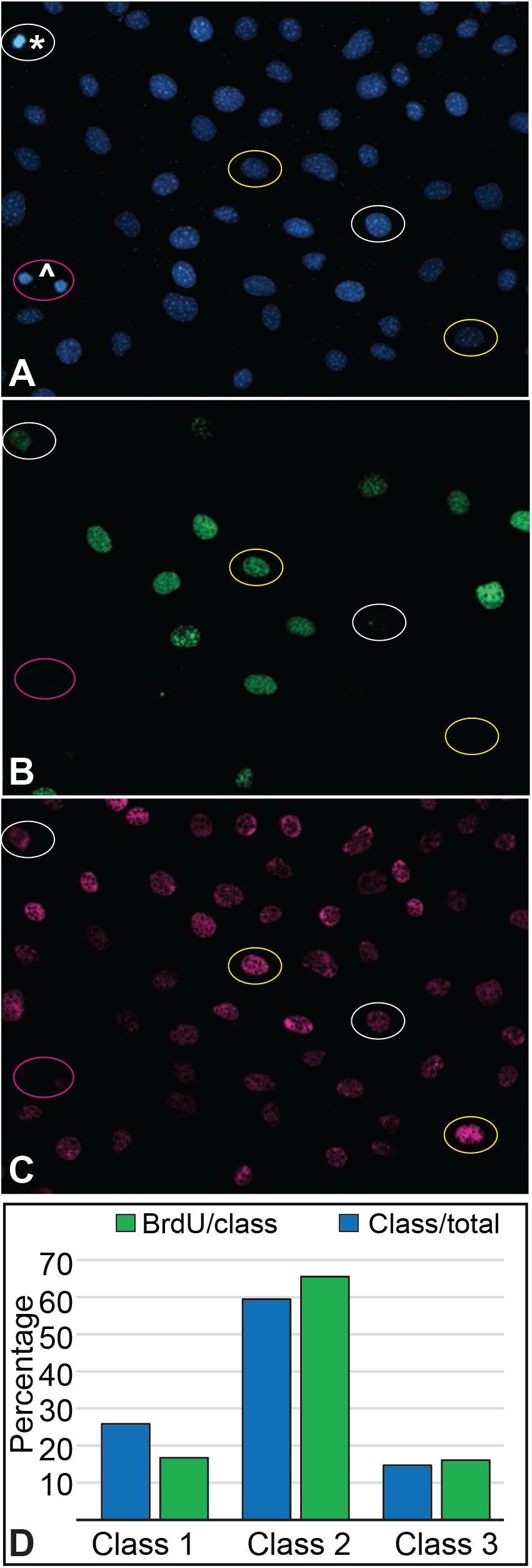
Satb2 levels exhibit population-level variation. Undifferentiated (T0) WT cells were stained with **A)** Hoechst to identify DNA (blue) and immuno-stained for **B)** BrdU (green) and **C)** Satb2 (pink). Three Satb2 expression classes were defined. Class 1 (pink ovals) represents low Satb2 levels, class 2 (white ovals) represents medium Satb2 levels, and class 3 (yellow ovals) represents high Satb2 levels. The asterisk indicates a metaphase cell, and the caret indicates recently divided, early G1 phase cells. **D)** Bar graph showing the percentage of total cells per Satb2 expression class (blue bars) and the percentage of each Satb2 class that was BrdU positive (green bars).

### 3.3 Mutations in Satb2 reduce pre-osteoblast proliferation

Previous work has reported a role for Satb2 in regulating osteogenic differentiation, but not found a role in osteoblast proliferation (Dobreva *et al.*, 2006). We find that mutations in one or more *Satb2* alleles result in reduced proliferative capacity of pre-osteoblasts (Fig. 4). All of the mutant cell lines proliferate more slowly than WT (Fig. 4A). Notably, colony 4, which expresses only Satb2 protein with the 40aa N-terminal deletion, proliferates even more slowly than all of the other lines, including colony 8, which lacks all Satb2 protein. Additionally, *Satb2* mutant cells have a greater than 50% reduction in mitotic cells compared to WT cell lines (Table 1).

**Table 1:** Quantification of nuclear morphology. Fixed cells were stained with Hoechst to visualize nuclear morphology. Nuclei were scored for mitosis, apoptosis, large size, donut-like morphology, and nuclear blebbing. For each colony, an area of larger than 3 mm^2^ was counted. Images used for counting are available at http://dx.doi.org/10.17632/6yfs85wyy6.1.

| **nuclear phenotype** | **WT** | **C20** | **C19** | **C9** | **C8** | **C4** |
| --- | --- | --- | --- | --- | --- | --- |
| mitotic | 51 | 50 | 16 | 18 | 11 | 23 |
| apoptotic | 2 | 8 | 4 | 14 | 9 | 10 |
| chromatin bridge | 15 | 4 | 24 | 15 | 6 | 5 |
| large nucleus | 1 | 6 | 3 | 3 | 8 | 7 |
| donut nucleus | 1 | 1 | 2 | 5 | 2 | 0 |
| nuclear blebbing | 4 | 1 | 18 | 34 | 9 | 30 |

**Figure 4:**
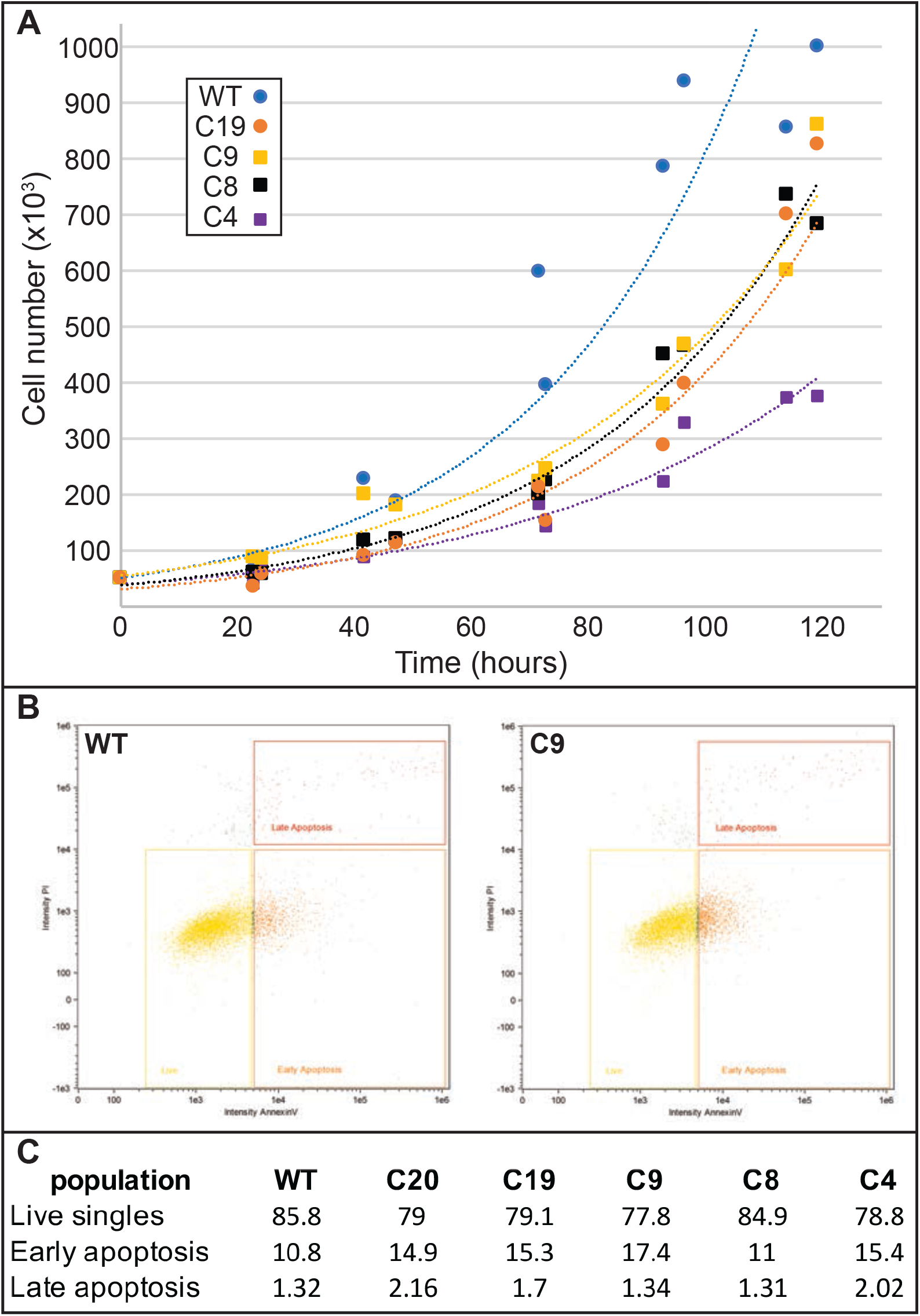
Mutations in *Satb2* reduce pre-osteoblast proliferation rates. **A)** Growth curves for undifferentiated (T0) cells. **B)** Representative flow cytometry detection of Annexin V (x-axis) and propidium iodide (y-axis) staining in WT (left) and C9 (right) cells. Yellow rectangles indicate live cells, orange boxes indicate early apoptotic cells, and red boxes indicate late apoptotic cells. **C)** Quantification of flow cytometry for all colonies is shown.

Decreased proliferation may be also be caused by increased apoptosis. We observed apoptotic cells in our immunostaining assays, but quantification of fixed cells did not indicate a significant increase in numbers (Table 1). This may be due to loss of cells during fixation and staining and/or lack of appropriate markers. Therefore, we turned to imaging flow cytometry to quantify dead and dying cells using Annexin V to identify early apoptotic and propidium iodide to identify late apoptotic cells. There is some variation in early apoptotic cell numbers, but very little difference in late apoptotic cell numbers (Fig. 4B,C). We observe minor variation in these percentages between experiments, but the overall pattern of only subtle increases in apoptosis in our mutant cell lines is consistent. Notably, our loss of function mutant (colony 8) has similar numbers of apoptotic cells as WT (and colony 20), yet still proliferates much more slowly. Taken together with our quantifications on fixed cells, these data indicate that mutations in *Satb2* mildly increase apoptosis, but that apoptosis is not sufficient to explain reductions in proliferation.

### 3.4 Mutations in Satb2 generate nuclear aberrations

In our immunostaining assays to detect Satb2 levels, we noticed a number of nuclear aberrations in the mutant cell lines. We defined 4 types of nuclear aberration (Fig. 5): chromatin bridges (Fig. 5A,B), large nuclei (Fig. 5C), nuclear blebbing or chromatin herniation (Fig. 5D-F), and donut-shaped nuclei (Fig. 5E,F). We quantified the abundance of these aberrations in fixed cells (Table 1). All 4 of these aberrations appear more frequently in all of our mutant cell lines relative to WT, suggesting that the aberrations are specific to mutations in *Satb2*. However, to address the possibility that these aberrations are a consequence of CRISPR/Cas9 treatment, we also quantified cells from colony 20, a line that was subjected to our CRISPR/Cas9 protocol but was genotyped as WT after selection.

**Figure 5:**
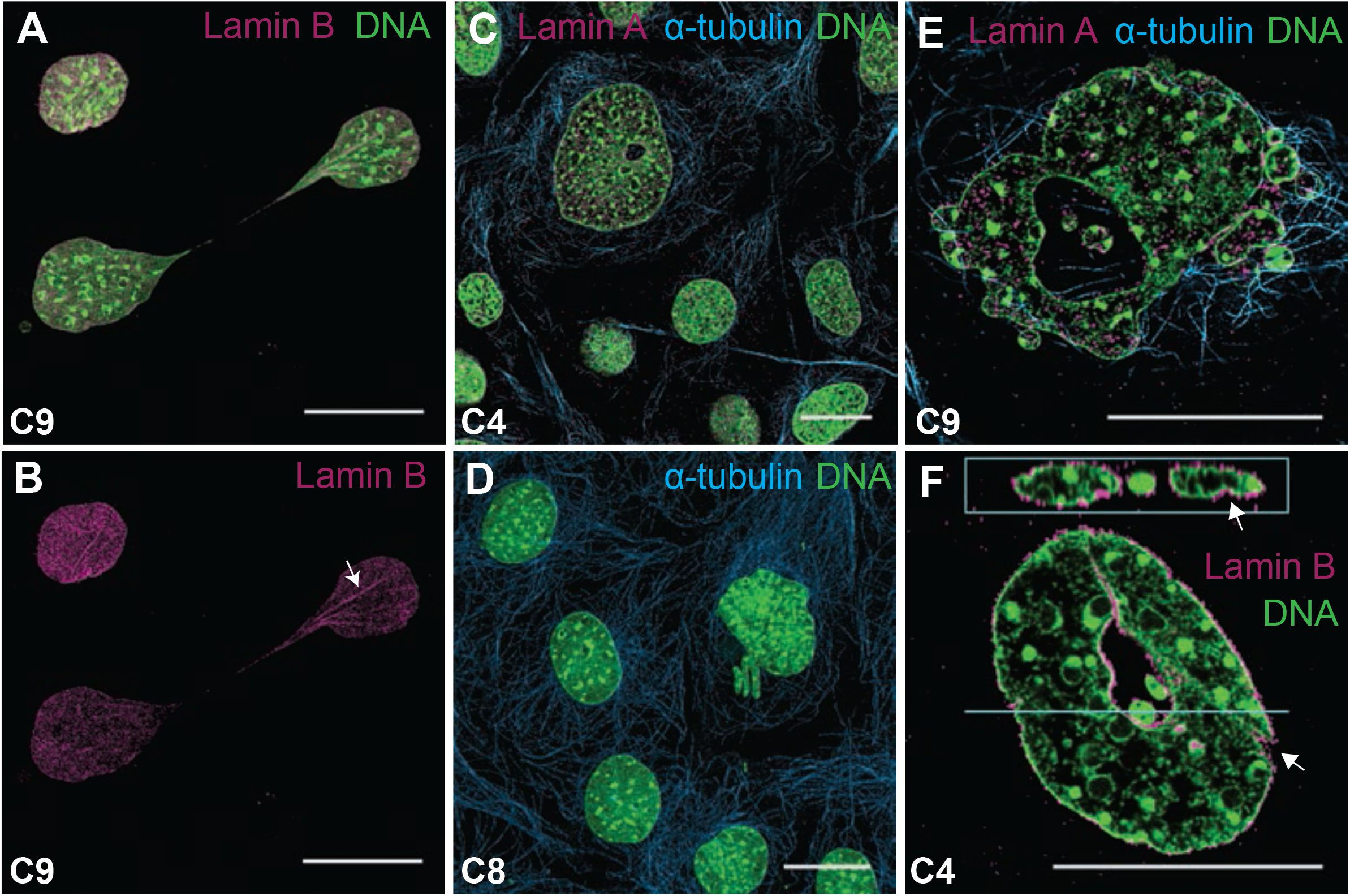
Mutations in Satb2 cause aberrant nuclear morphology. Confocal images of *Satb2* mutant osteoblasts showing representative nuclear aberrations. **A**,**B**) C9 cells with a chromatin bridge. Lamin B immunostaining highlights nuclear folds being pulled into bridge (arrow in B). **C**) Large cell with small hole shown relative to other normal-sized cells. **D**) Chromatin herniation from nucleus. **E, F**) Nuclear blebbing and donut-shaped nuclei in C9 (E) and C4 (F) cells. Note the presence of nuclear folds in F (arrows). DNA in green; alpha-tubulin in teal; lamin in pink (Lamin A in C and E; Lamin B in A, B, and F). Images in A, B, C, and D are maximum intensity projections. Images in E and F are single confocal sections. Panel F includes a z-section at the top. All scale bars represent 10 µm.

In both WT and colony 20 cells, the presence of nuclear aberrations was rare, and always more subtle than those observed in the mutant cell lines. For example, chromatin bridges, which was the most common aberrant phenotype seen in WT cells, are typically present as very thin string-like strands between nuclei (see Fig. 2B). In contrast, chromatin bridges in the mutant cell lines were often thick as shown in Figure 5B, where Lamin B immunostaining shows folds in DNA within the bridge. Similar folds in DNA can be observed in Figure 5F. Finally, while WT cells exhibit the occasional small nuclear bleb, we only observe the catastrophic blebbing shown in Figure 5E in the mutant cell lines. Large panel overviews of all 6 cell lines from which quantifications were made are available on Mendeley (http://dx.doi.org/10.17632/6yfs85wyy6.1).

### 3.5 Mutations in Satb2 reduce osteogenic differentiation

To investigate how *Satb2* mutations in our cell lines affect osteogenic differentiation, we performed an *in vitro* differentiation assay (Fig. 6). Differentiation was evaluated by histology (alizarin red) and gene expression (qPCR). By both measures, all of our mutant cell lines, even colony 19, a true heterozygote, exhibit incomplete differentiation relative to WT cells (Fig. 6). We do observe mineralization, as reflected by alizarin red staining, in all of our colonies. However, none of the colonies form as many nodules of punctate staining as observed in WT cells (Fig. 6B). Additionally, the expression of bone matrix genes is much lower in all *Satb2* mutant cell lines (Fig. 6C).

**Figure 6:**
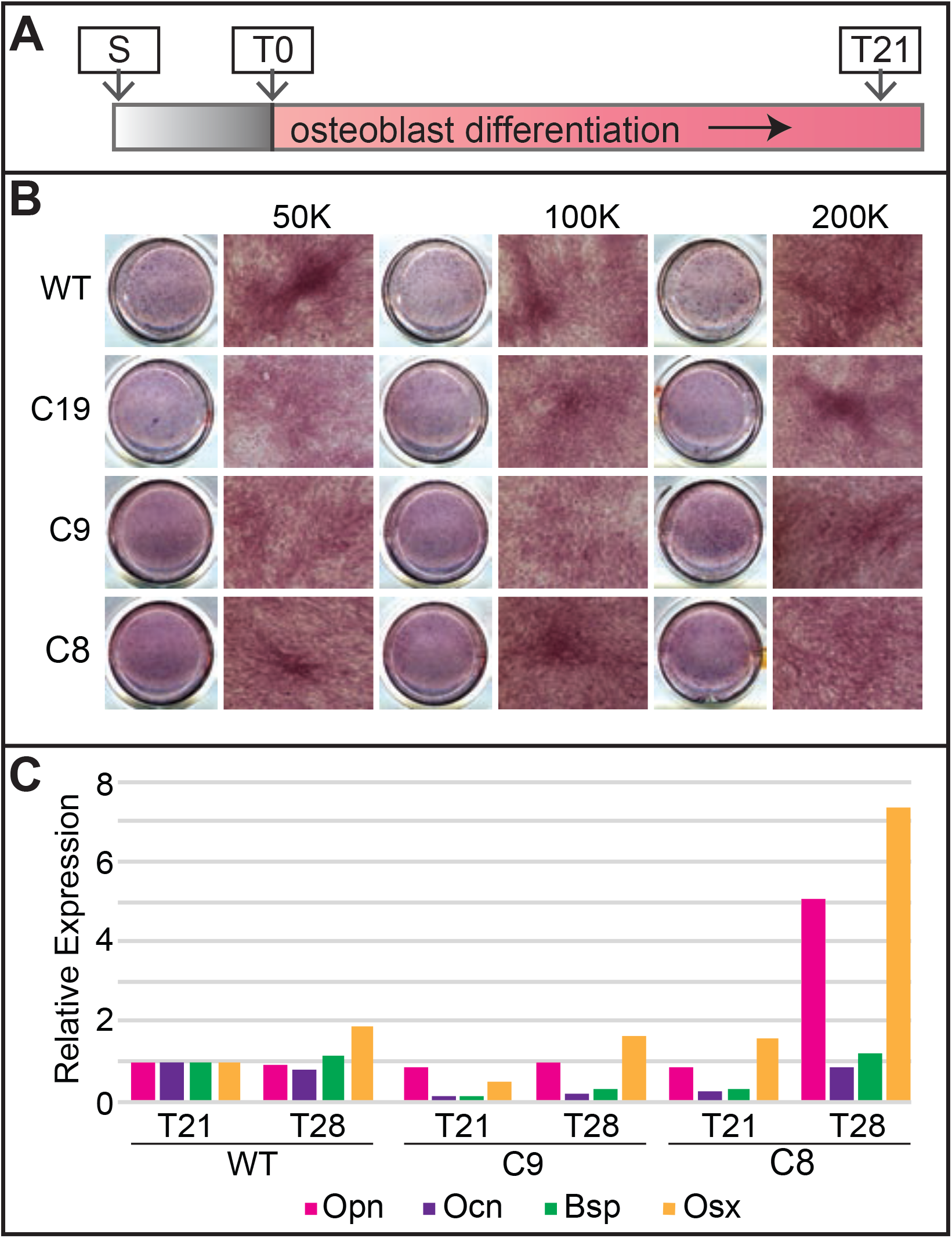
Mutations in Satb2 reduce, but don’t inhibit, osteogenic differentiation. **A)** In vitro differentiation assay timeline. **B)** Overview scans of wells (left) and 10x magnification of Alizarin Red staining on WT, C19, C9, and C8 cells after 21 days (T21) in differentiation. Data are shown for seeding densities of 50,000 cells, 100,000 cells, and 200,000 cells. Note that clusters start to form when mutant cells are plated at higher density (compare asterisk in C19 at 200K to WT at 50K). **C)** Expression of *Opn* (pink bars), *Ocn* (purple bars), *Bsp* (green bars), and *Osx* (yellow bars) is shown relative to WT at T21.

As reported above, our mutant cell lines have proliferation defects. Therefore, we tested whether differentiation potential was affected by cell density at the onset of differentiation. We plated cells at 3 different densities prior to differentiation, and also extended our differentiation protocol by an additional week (28 vs. 21 days). Increasing cell number at the onset of differentiation appears to improve osteogenic output as determined by the presence of mineralized nodules (Fig. 3B). Additionally, extending the differentiation time leads to increased expression of bone matrix proteins (Fig. 3C). Interestingly, colony 8, the loss of function line, differentiates better than colonies 9 and 19, two heterozygous lines. These data suggest that proliferation defects reducing cell density at the onset of differentiation contribute to reduced osteogenesis in cells with *Satb2* mutations.

### 3.6 Satb2 mutations increase gene expression variance

We used single cell gene expression analyses to evaluate how mutations in *Satb2* affect osteogenic gene expression. Expression was quantified for 93 genes including 66 related to osteogenesis, 18 related to cell cycle progression, and 7 related to apoptosis (**Supplementary Table 1**). Down-regulation of *Satb2* alters the expression profile of differentiating (T14) osteoblasts (Fig. 7 and **SupFig. 3**). Single cell expression profiles can be visualized using tSNE plots (Fig. 7A,B) which represent all the variation in the dataset and PCA plots (Fig. 7C,D) which represent 46% and 28% of the variation, respectively. As differentiation proceeds, gene expression becomes increasing distinct between WT and mutant cells. At T0, there is no significant difference between the means or variance the two populations, however by T14, gene expression shows more variance between individual mutant cells than is observed in WT cells (T0, F=1.11, n.s; T14, F=3.19, p=0.01).

**Figure 7:**
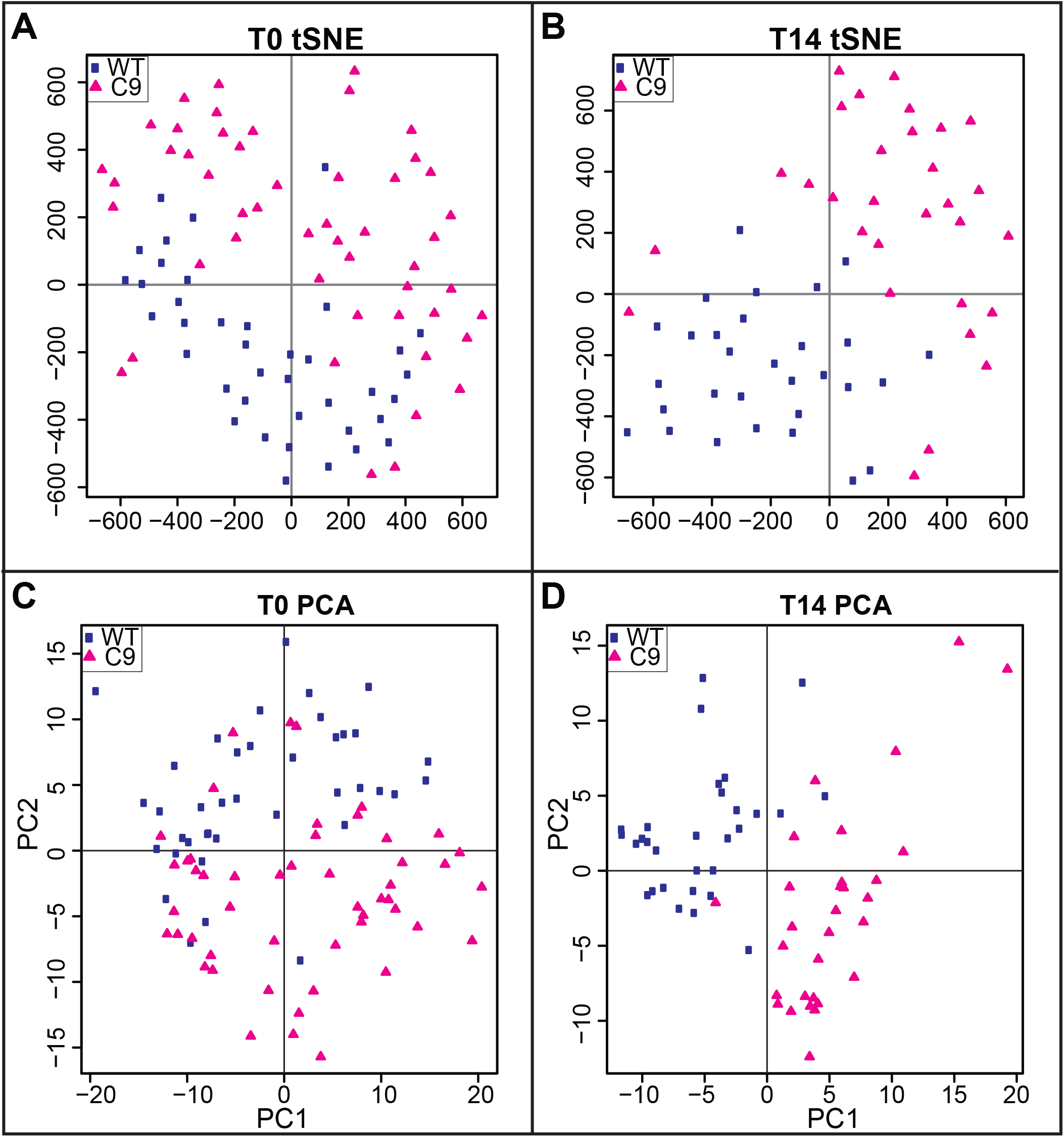
Reduction in Satb2 increases gene expression variance. Upper panels show t-distributed Stochastic Neighbor Embedding (tSNE) plots of single cell gene expression in wild-type (WT; blue squares) and colony 9 (C9; pink triangles) cells at **A)** T0 (initial dimensions= 80, perplexity=15, iterations 4000, error was 0.528 at the 4000th iteration) and **B)** T14 (initial dimensions= 80, perplexity=10, iterations 3000, error was 0.524 at the 3000th iteration). Lower panels show plots of principal components 1 and 2 from Principal Components Analysis (PCA) on single cell gene expression in WT (blue squares) and C9 (pink triangles) cells at **C)** T0 and **D)** T14.

The increased variation of gene expression patterns in mutant cells can also be observed by evaluating the number of genes that correlate with *Satb2* (**SupFig. 3**). At T14, the expression of 16 genes positively correlates with *Satb2* in WT cells (correlation coefficient of 0.5 and above). In contrast, only 4 genes show correlated expression with *Satb2* in T14 mutant cells. No genes in either cell population negatively correlate with *Satb2*. These data suggest that large-scale alterations to gene expression occur when *Satb2* is down-regulated, and that *Satb2* mutations lead to increased population variation in osteoblast gene expression.

## 4. Discussion

### 4.1 Cell biological mechanism of osteogenic defect

Although it is generally accepted that loss of Satb2 causes bone hypoplasia and low bone mineral density, there is some discrepancy among the reported molecular and cellular mechanisms mediating these phenotypes. In a study focusing on the developing jaw, loss of *Satb2* was associated with progenitor cell death, and more specific effects of Satb2 on osteogenic differentiation (in the facial or axial skeleton) were not reported (Britanova *et al.*, 2006). That study also found that *Satb2*^+/-^ heterozygous mice have phenotypes of the craniofacial skeleton that are intermediate between WT and mutant. In a separate study, loss of *Satb2* was associated with deficiencies in osteogenic gene expression (Dobreva *et al.*, 2006). Notably, Dobreva and colleagues (2006) reported that *Satb2*^+/-^ mice were phenotypically normal and used them as controls to evaluate gene expression defects in *Satb2*^-/-^ mice. In addition to not finding evidence of apoptosis (even in homozygous mutants), Dobreva and colleagues (2006) also report that proliferation of osteoblasts was not affected, although these data were not shown. In general, the craniofacial phenotype shown by Dobreva and colleagues (2006) appears less severe than that shown by Britanova and colleagues (2006). Here, we find that mutations in *Satb2* cause both proliferation and differentiation defects. We also find a dosage effect, where loss of even a single allele of *Satb2* causes defects in osteogenesis. Thus, we hypothesize that Satb2 has two cell biological roles in osteogenesis: 1) regulation of pre-osteoblast proliferation and 2) subsequent osteoblast differentiation (Fig. 8).

**Figure 8:**
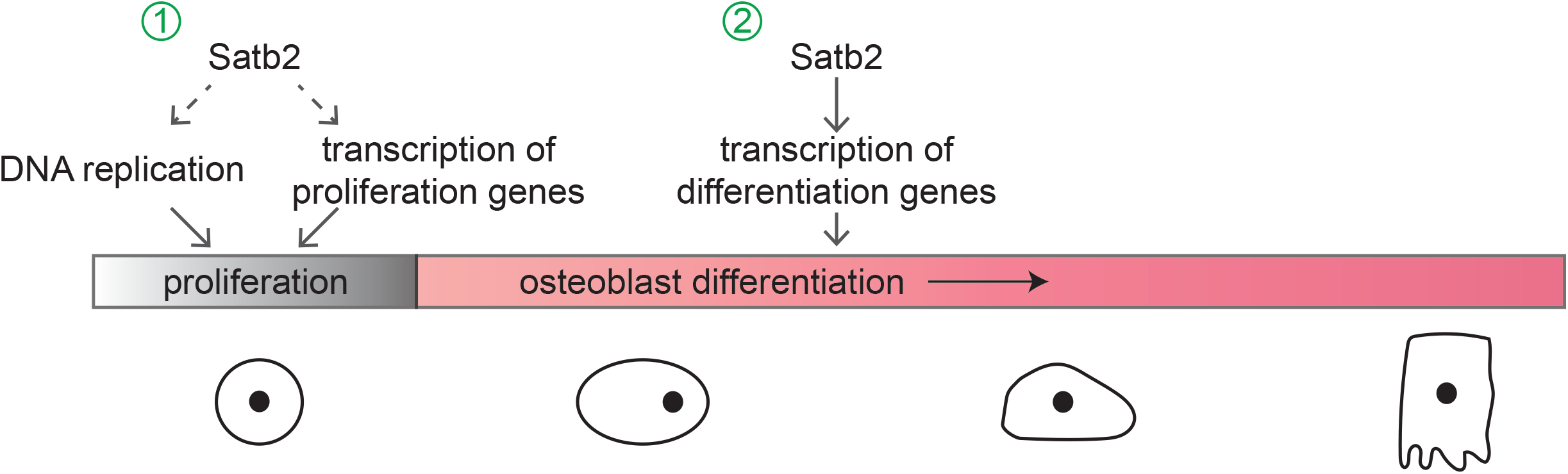
Satb2 has two distinct functions during osteogenesis. Osteogenic differentiation time-line showing 1) an early role for Satb2 mediating pre-osteoblast proliferation, and 2) a later role during differentiation.

The different results reported in the previous studies described above may be related to differences in genetic background (although the background used in the Dobreva study was not specified). Our work, conducted in the C57BL/6 background (which is the same background used by Britanova and colleagues), indicates that *Satb2* mutations increase apoptosis (although not to the extent they observed) in pre-osteoblasts. Additionally, we show that increasing cell density at the onset of proliferation increases differentiation potential. Taken together, these data indicate that proliferation defects in osteogenic progenitors contribute to deficits in bone mineral production, particularly in the C57BL/6 background. Disruption to osteogenic gene expression has been reported in all loss of *Satb2* studies, suggesting it is a more penetrant phenotype. If the C57BL/6 background is susceptible to both proliferation and differentiation defects, it would also explain why the jaw phenotype reported by Britanova and colleagues (2006) is more severe. Further, this data raises the interesting possibility for modifiers with the potential to buffer the proliferation defects to be discovered.

### 4.2 Molecular mechanism of osteogenic defect

MAR-binding proteins have at least two distinct molecular roles, transcriptional regulation and DNA replication (Ottaviani *et al.*, 2008b). MAR-binding proteins regulate transcription by defining the borders of chromatin loops anchored in the nuclear matrix. The anchoring sites form discrete territories of genome organization that serve as transcription hubs, where transcription factors, histone modifying enzymes, and other proteins aggregate to promote gene expression (Cai *et al.*, 2003; Gyorgy *et al.*, 2008; Kirillov *et al.*, 1996; Ottaviani *et al.*, 2008b). Mutations reducing MAR-binding proteins may limit their ability to properly modify chromatin and/or participate in protein-protein interactions mediating gene transcription thus disrupting the coordinated regulation of cell-type specific genes. For example, in *Satb1* mutant mice, gene expression is dysregulated during differentiation of thymocytes and T cells (Alvarez *et al.*, 2000). Similarly, we observe a significant shift in gene expression and increased variation in gene expression in *Satb2* mutant cells. We do not observed increased variance in gene expression in proliferating osteoblasts (T0), but this may be an artifact of the genes used in our single-cell analysis, which are heavily biased towards osteogenic differentiation genes.

However, our finding that proliferation is affected in pre-osteoblasts indicates an additional role for Satb2. Satb2 may also mediate this effect through regulation of transcription of genes important for cell cycle progression. Notably, in a different study in MC3T3-E1 cells where *Satb2* was over-expressed, genes associated with cell cycle and proliferation were more upregulated than genes associated with differentiation (Kim *et al.*, 2012). However, the possibility that Satb2 is more directly involved in DNA replication by binding at MARs remains open, especially as Satb2 is highly expressed from late G1 through G2. If Satb2 is required for DNA replication (either directly through binding at MARs or indirectly through gene regulation), its reduction may cause replication stress, which is associated with DNA damage (Zeman and Cimprich, 2014). Further investigation into Satb2 involvement in DNA replication could elucidate mechanisms underlying DNA damage observed in *Satb2* mutants.

### 4.3 N-terminal domain of Satb2

Several genetic modifications of the *SATB2* locus, including deletions, point mutations, duplications, chromosomal translocations, and frameshift and splice site mutations have been implicated in causing SAS (Zarate and Fish, 2017). Most of these mutations have been hypothesized to mediate disease phenotypes through haploinsufficiency of *SATB2*. However, few mutations have been functionally tested. In the case of one nonsense mutation, the mutant *SATB2* mRNA produces a truncated protein retaining the SATB2 dimerization domain, and lacking the DNA-binding motifs, thus suggesting a dominant negative effect (Leoyklang *et al.*, 2007; Leoyklang *et al.*, 2013). Some SATB2 activity remains in cells expressing this truncated protein, however, it remains unclear how these levels compare to *SATB2* mutations resulting in loss of function. More generally, it is still not well understood how heterozygous loss of function mutations affect *SATB2* mRNA and protein levels.

Here we show that mutations in Satb2 result in decreased mRNA and protein levels. Additionally, several observations from our work suggest that mutations may generate proteins with dominant negative effects. First, colony 4 (generating only protein with the N-terminal deletion) proliferates more slowly than colony 8 (complete loss of function). Colony 8 appears to have fewer nuclear aberrations than either colony carrying the N-terminal deletion. Finally, colony 8 differentiates better than the colonies carrying the N-terminal deletion. We used the Phyre^2^ software to predict the effect of this deletion on Satb2 structure (Kelley *et al.*, 2015). Although we did not see any obvious alterations to Satb2 protein folding, we did find that the N-terminal domain is highly disordered (**SupFig. 4**). Disorder domains are important for protein dissociation (Umezawa *et al.*, 2016). Chromatin architecture is dynamic and genomic anchors need to be altered throughout the cell cycle to adapt to changing genomic function (Heng *et al.*, 2004; Ottaviani *et al.*, 2008a). Additionally, chromatin remodeling complexes dissociate from chromatin during mitosis (Ma *et al.*, 2015). Mutations in the Satb2 N-terminal region may therefore hinder its ability to dissociate from chromatin, disrupting its function and/or contribute to DNA damage.

### 4.5 Variation in disease phenotypes

Individuals with SAS exhibit a broad spectrum of skeletal abnormalities including tibial bowing, osteomalacia, osteopenia or osteoporosis, and they have a high risk for fractures (Zarate *et al.*, 2018b). The incidence and severity of these skeletal anomalies varies among SAS patients, with some individuals having little to no skeletal problems (Zarate *et al.*, 2018a). Age at presentation also reflects variability of severity with skeletal morbidity reported as early as infancy. Current available data suggests that for those individuals that have undergone dedicated evaluation for bone density, it is common to have lower age/gender adjusted BMD scores regardless of the type of genetic modification in *SATB2*. Overall, the current data from SAS patients is consistent with variation in penetrance of skeletal anomalies that is not associated with mutation type. We hypothesize that genetic background underlies these differences, with more severe phenotypes occurring in individuals that are more susceptible to pre-osteoblast proliferation defects.

Anti-resorptive drugs (e.g. bisphosphonates) which increase bone density and strength by inhibiting osteoclast activity, have been the main therapeutic option for individuals with childhood osteopenia in SAS (Zarate *et al.*, 2018b). The results presented here showing a role for *Satb2* in pre-osteoblast proliferation suggest that the low BMD in SAS may be a consequence of a decline in the number and vitality of osteoblasts. Therefore, drugs that directly target osteoblasts such as teriparatide, an anabolic agent that increases osteoblast formation and activates bone-lining cells (quiescent osteoblasts) to form new bone, could provide a biologically sound alternative treatment for osteopenia in SAS.

### 4.6 Implications for cancer

We observed several types of nuclear aberrations, including nuclear blebbing and chromatin herniation, that are also characteristic of cancer cells. A previous study in MC3T3-E1 cells found that *Satb2* levels increase in response to oxidative stress. When *Satb2* was reduced via siRNA in cells under oxidative stress, apoptosis increased (Wei *et al.*, 2012). Similarly, Satb1 was recently found to participate in DNA repair associated with oxidative damage (Kaur *et al.*, 2016). Reductions in *Satb2* have also been reported in cancers associated with deficiencies in DNA mismatch repair proteins (Ma *et al.*, 2018). Together, these data suggest that down-regulation of Satb2 may be involved in cancer through compromised DNA repair mechanisms. Paradoxically, over-expression of Satb2 is also associated with cancer in different cell types (Chen and Costa, 2018; Ma *et al.*, 2018). Our finding that Satb2 is involved with proliferation and DNA damage may explain this dichotomy. Cell-type specific gene regulation may impact how alterations to Satb2 disrupt normal cell biology in cancer progression.

## 5. Conclusion

Previous work has described Satb2 is a high-order transcription factor regulating genes required for osteogenic differentiation. Our data suggest that, in addition to its role in differentiation, Satb2 regulates progenitor proliferation. Satb2 may promote pre-osteoblast proliferation through regulation of the expression of genes associated with cell cycle progression and/or it may have a role during DNA replication through its association with MARs. The involvement of Satb2 in separate molecular processes during osteogenesis, one of which may be buffered in certain genetic backgrounds, may help explain variation in disease phenotypes (Merkuri and Fish, 2019). We have also found that mutations in *Satb2* cause chromatin defects. Further investigation of the mechanism underlying these defects has potential impacts on both skeletal defects and cancer.

## Contributions

TD, EES, JD, and JLF designed and conducted experiments. TD, EES, JD, FM, YZ, and JLF analyzed the data. EES, YZ, and JLF wrote the manuscript. All authors agree on the results and commented on the manuscript.

## Acknowledgements

This work was funded by NIH R15DE026611. Generous technical help was provided by Jack Lepine (UML), Susanne Pechold (UMMS) and Kahraman Tanriverdi (UMMS). We thank the SATB2 Foundation and SAS families for sharing their stories and providing inspiration to solve mysteries underlying this disease.

