## Supplemental Figures for "Satb2 regulates proliferation and nuclear integrity of pre-osteoblasts"

#### Supplemental Data

**Sup Fig. 1**: Mutations in *Satb2* reduce protein levels
**A)** Satb2 expression determined by qPCR. Green bars reflect expression detected with primers binding at the mutation site. Note the absence of expression for these primers in C9, C4, and C8, all colonies lacking the N-terminal region of Satb2. C19 has one WT allele and produces roughly 60% of WT levels of Satb2 full-length mRNA. Orange bars reflect expression detected with primers binding in the C-terminal region of Satb2. Note that all colonies produce mRNA, although not all of this RNA is expected to make protein. **B)** Western Blot using an antibody directed to the C-terminal region of Satb2. Note that C19 produces roughly half the amount of Satb2 protein as WT, as predicted by mRNA levels. C9 produces two separate proteins, including a shorter protein predicted to lack to N-terminal 40 amino acids (asterisk). C4 also produces the short protein although at much lower levels, and C8 produces no protein.

**SupFig2:** *Satb2* expression is more closely related to cell cycle genes than osteogenic genes
Heatmap comparing single cell gene expression. Individual cells are in columns. Genes are in rows. A cluster of highly expressing *Satb2* cells (black rectangle) is associated with high levels of cell cycle regulators associated with the S and G2 phases of the cell cycle (bold green). *Satb2* is listed in bold red.

**Sup Fig 3:** Mutations in *Satb2* dysregulate gene expression during differentiation
Genes whose expression is positively correlated with Satb2 with a correlation coefficient 0.5 and above are shown for **A)** WT and **B)** C9. Each point on the x-axis shows expression levels in a single cell (30 cells for WT; 26 cells for C9).

**Sup Fig 4:** Satb2 N-terminal disorder domain and protein folding
Predicted structure of **A)** wild-type (WT) and **B)** 40 amino acid N-terminal deletion (MT) proteins produced by the Phyre$^2$ software (Kelley *et al.*, 2015). **C)** Sequence, secondary structure, and disorder confidence predictions for the 40 amino acid N-terminal region of Satb2.

**Supplementary Table 1:** List of Fluidigm Delta Gene Assays
Genes and primer sequences used for single-cell qPCR Biomark analysis are listed.

### Supplemental Figure 1

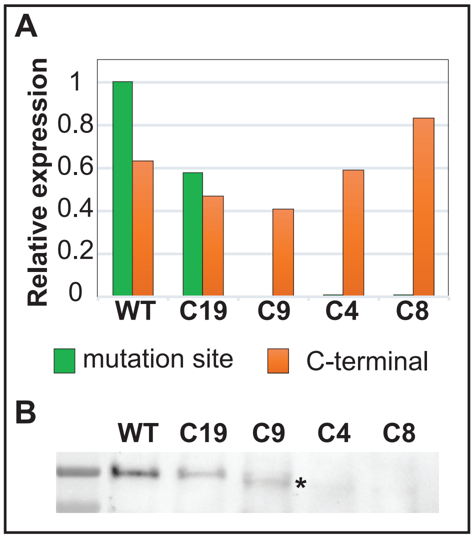

#### Supplemental Figure 2

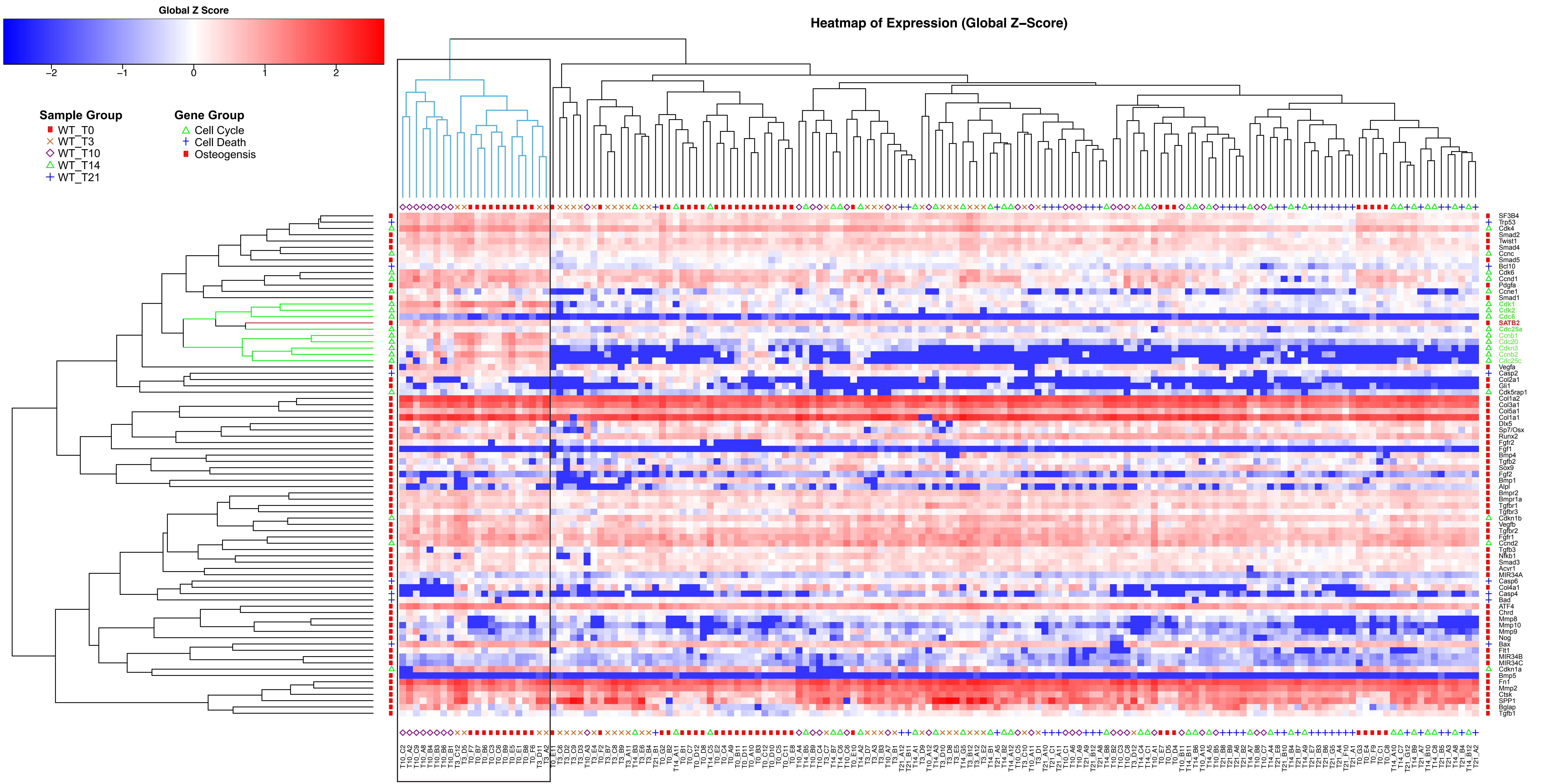

### Supplemental Figure 3

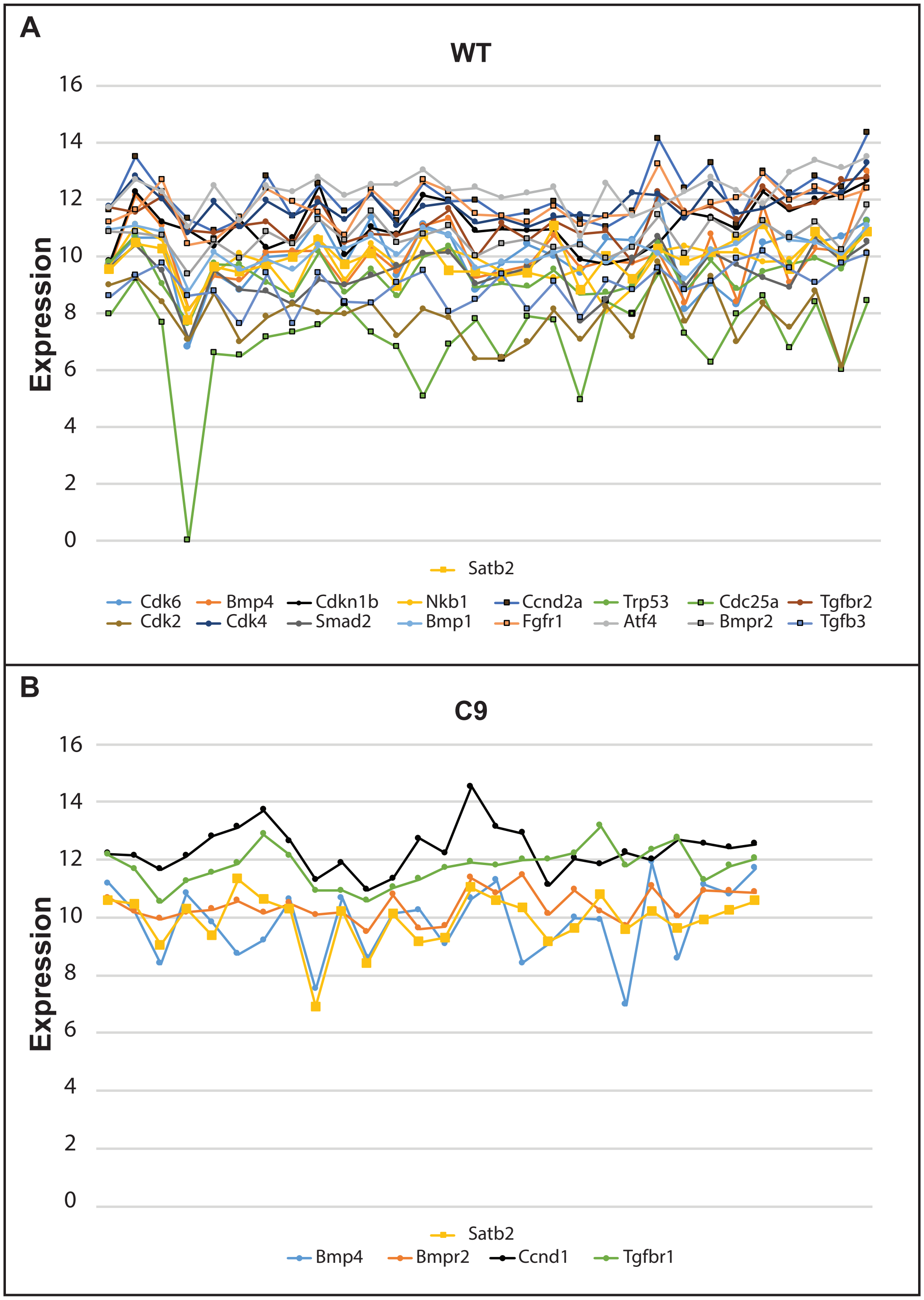

### Supplemental Figure 4

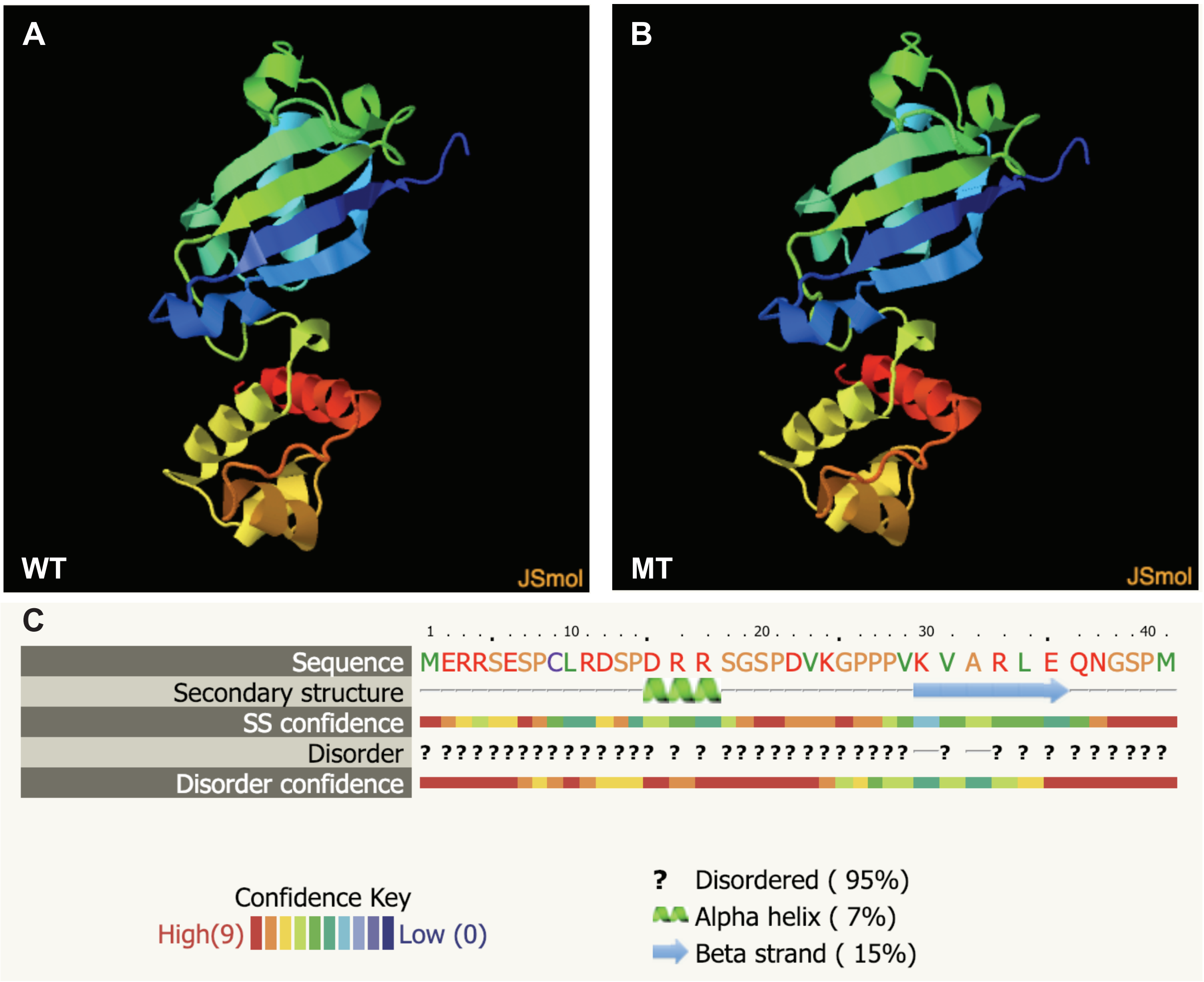

| Target | Forward Primer | Reverse Primer | Design RefSeq | Gene Symbol |
|---|---|---|---|---|
| **Actb** | ACAGCTTCTTTGCAGCTCCT | AGCGCAGCGATATCGTCAT | NM_007393.5 | Actb |
| **Acvr1** | TGATGATGGCTTTCCCTTCC | CCCTCACACACACACATGTA | NM_007394.3 | Acvr1 |
| **Alpl** | GCAATGAGGTCACATCCATCC | GTTCACCCGAGTGGTAGTCA | NM_007431.2 | Alpl |
| **ATF4** | GCCAAGCACTTGAAACCTCA | CATCCAATCTGTCCCGGAAAA | NM_009716.2 | Atf4 |
| **Bad** | GCAGCCACCAACAGTCATCA | GTACGAACTGTGGCGACTCC | NM_007522.2 | Bad |
| **Bax** | GCGTGGTTGCCCTCTTCTA | CTGATCAGCTCGGGCACTTTA | NM_007527.3 | Bax |
| **Bcl10** | CTCACGGAGGAGGATTTGAC | TTTCTCACACAGGTAAACACGTA | NM_009740.1 | Bcl10 |
| **Bglap** | GAACAGACAAGTCCCACACA | GTCAGCAGAGTGAGCAGAAA | NM_007541.2 | Bglap |
| **Bmp1** | CCGTTTGTGATTGGAGGGAA | ATGCTTCTCCCAGTGTCTCA | NM_009755.3 | Bmp1 |
| **Bmp2** | GTGCGCAGCTTCCATCAC | CGTCACTGGGGACAGAACTTAA | NM_007553.2 | Bmp2 |
| **Bmp3** | CGGCTGGAGCGAATGGATTA | GTGGCGTGATTTGATGGTTTCA | NM_173404.3 | Bmp3 |
| **Bmp4** | GAACCGGGCTTGAGTACCC | GGTCCCTGGGATGTTCTCC | NM_007554.2 | Bmp4 |
| **Bmp5** | CTGCAAGAAGCACGAACTCTA | AGCATACCCTTCTGGTGCTA | NM_007555.3 | Bmp5 |
| **Bmp6** | CCAAGTCTTGCAGGAGCATCA | CCTTCTTCTGAGGCCCACAC | NM_007556.2 | Bmp6 |
| **Bmp7** | TGTACGTCAGCTTCCGAGAC | GTAGTAGGCAGCATAGCCTTCA | NM_007557.2 | Bmp7 |
| **Bmpr1a** | GGAAATGGCTCGTCGTTGTA | CACTGGGCACCATGTTGTAA | NM_009758.4 | Bmpr1a |
| **Bmpr1b** | GAAGTTACGGCCTTCATTCCC | GCACTCTGTCATAAGCTTCCC | NM_007560.3 | Bmpr1b |
| **Bmpr2** | GGGGAAGAAGATAATGCGGCTA | GGTTCACAGCTCCTTCTAGCA | NM_007561.3 | Bmpr2 |
| **Casp2** | TCCAAGTCTACAGAACAAGCCAAA | TCCATCTTGCTGGTCGACAC | NM_007610.1 | Casp2 |
| **Casp4** | CCACGTGGAGAAGGACTTCA | TCCTGTTTTGTCTCGGTAGGAC | NM_007609.3 | Casp4 |
| **Casp6** | ACAGAAACCGATGGCTTCTACA | TCCTCTTGTGGTCCATCTTGTAC | NM_009811.4 | Casp6 |
| **Ccnb1** | GCTGCTTCAGGAGACCATGTA | TAGCATCTTCTTGGGCACACA | NM_172301.3 | Ccnb1 |
| **Ccnb2** | GCCTCTTGCCTGTCTCAGAA | CACTCTCCATGTAGCCTGTGTA | NM_007630.2 | Ccnb2 |
| **Ccnc** | ACGGATCTCTGTCTGCTGTA | TGTTGTACGACACAGGCTAC | NM_016746.3 | Ccnc |
| **Ccnd1** | TGCCGAGAAGTTGTGCATCTA | TGTTCACCAGAAGCAGTTCCA | NM_007631.2 | Ccnd1 |
| **Ccnd2** | TGCGGAAAAGCTGTGCATTTA | ACCCAACACTACCAGTTCCC | NM_009829.3 | Ccnd2 |
| **Ccne1** | CTGTGAAAAGCGAGGATAGCA | TGTTAGGGGTGGGGATGAAA | NM_007633.2 | Ccne1 |
| **Cdc20** | TCCCCTGCAAACATTCACTCA | ATATTGGACTGCCAGGGACAC | NM_023223.2 | Cdc20 |
| **Cdc25a** | TTGGACAGTGACCCAAGAGAC | AATCCTGATGCTTCCCAGAGAC | NM_007658.3 | Cdc25a |
| **Cdc25c** | GATGTCTGCCTCTCCGCTTA | CTATCCAGAGGTCCAGATGAATCC | NM_009860.2 | Cdc25c |
| **Cdc6** | TCTGCTGTTTCAGGAGACATCC | CCTGACATCCGACTCCACAA | NM_001025779.1 | Cdc6 |
| **Cdk1** | AAGTACCTGGACTCCATCCC | TCCCTGGAGGATTTGGTGTAA | NM_007659.3 | Cdk1 |
| **Cdk2** | GGAGAAGTTGTGGCGCTTAA | GAGATCTCTCGGATGGCAGTA | NM_183417.3 | Cdk2 |
| **Cdk4** | ATGTGGAGCGTTGGCTGTA | TGGTCGGCTTCAGAGTTTCC | NM_009870.3 | Cdk4 |
| **Cdk5rap1** | TCCTCACCCCAAGGATTTTCC | CTGGCAGGTGGATCTGCTTA | NM_025876.2 | Cdk5rap1 |
| **Cdk6** | TGGAAGTTCAGACGTGGATCA | GTCCCTAGGCCAGTCTTCC | NM_009873.2 | Cdk6 |
| **Cdkn1a** | GAACATCTCAGGGCCGAAAAC | TCTGCGCTTGGAGTGATAGAA | NM_007669.4 | Cdkn1a |

| | | | | |
|---|---|---|---|---|
| **Cdkn1b** | CAGTGTCCAGGGATGAGGAA | TTCGGGGAACCGTCTGAAA | NM_009875.4 | Cdkn1b |
| **Cdkn3** | CAGACGAAGAACCTGTTGATGAA | ATTCACTCGCGACAGAGGTA | NM_028222.1 | Cdkn3 |
| **Chrd** | CCTTTGGGGAGATGAGCTGTA | GAACAATCGTCCCGCTCAC | NM_009893.2 | Chrd |
| **Col10a1** | TTGCTAGCCCCAAGACACAA | GCCTTGTTCTCCTCTTACTGGAA | NM_009925.4 | Col10a1 |
| **Col14a1** | TGGTCAGATCAAGAGGACCAA | GGCCCATGATGTAGAGCAAA | NM_181277.3 | Col14a1 |
| **Col1a1** | TTCAGGGAATGCCTGGTGAA | ACCTTTGGGACCAGCATCA | NM_007742.3 | Col1a1 |
| **Col1a2** | GAAAAGGGTCCCTCTGGAGAA | AATACCGGGAGCACCAAGAA | NM_007743.2 | Col1a2 |
| **Col2a1** | GCACTTGCCAAGACCTGAAA | CCTGGTTGGGATCAATCCAGTA | NM_031163.3 | Col2a1 |
| **Col3a1** | TGCTGGAAAGAATGGGGAGAC | GGTCCAGAATCTCCCTTGTCAC | NM_009930.2 | Col3a1 |
| **Col4a1** | TCTGGCTGTGGAAAATGTGAC | TCCAATGACACCTTGCAACC | NM_009931.2 | Col4a1 |
| **Col5a1** | GGTCCTTTGGGGAAACCA | CTGGAGGACCTTCTTTTCCA | NM_015734.2 | Col5a1 |
| **Ctsk** | AGGGAAGCAAGCACTGGATA | TTCCGAGCCAAGAGAGCATA | NM_007802.3 | Ctsk |
| **Dlx5** | TCTCTAGGACTGACGCAAAC | TGACTGTGGCGAGTTACAC | NM_010056.2 | Dlx5 |
| **Fgf1** | TGGACACCGAAGGGCTTTTA | GCATGCTTCTTGGAGGTGTAA | NM_010197.3 | Fgf1 |
| **Fgf2** | TCTTCCTGCGCATCCATCC | GCACACACTCCCTTGATAGACA | NM_008006.2 | Fgf2 |
| **Fgfr1** | GAGTAAGATCGGGCCAGACA | TCCATTTCCTTGTCGGTGGTA | NM_001079908.1 | Fgfr1 |
| **Fgfr2** | TCAAGTGGATGGCTCCTGAA | CACATTAACACCCCGAAGGAC | NM_010207.2 | Fgfr2 |
| **Flt1** | TTGCACGGGAGAGACTGAAA | GCCAAATGCAGAGGCTTGAA | NM_010228.3 | Flt1 |
| **Fn1** | GGAACCAGCAGAGTCCCAAA | CCTCGGTGTTGTAAGGTGGAA | NM_001276413.1 | Fn1 |
| **Gapdh** | CAAGGTCATCCCAGAGCTGAA | CAGATCCACGACGGACACA | NM_008084.2 | Gapdh |
| **Gdf10** | CCCAAATCCTTTGACGCCTAC | CAATGCCCACAGCTCTGAC | NM_145741.2 | Gdf10 |
| **Gli1** | CAGAATCGGACCCACTCCAA | GCGAGCTGGGATCTGTGTA | NM_010296.2 | Gli1 |
| **HOXA2** | GAAGAAGGCGGCCAAGAAA | GCTGCCATCAGCTATTTCCA | NM_010451.1 | Hoxa2 |
| **Ihh** | TCTTCAAGGACGAGGAGAACAC | AGATGGCCAGTGAGTTCAGAC | NM_010544.2 | Ihh |
| **MIR34A** | TGTCTTAGCTGGTTGTTGTGAGTA | GCAGCACTTCTAGGGCAGTA | NR_029751.1 | Mir34a |
| **MIR34B** | CTCGGTTTGTAGGCAGTGTAA | TGTTTTGATGGCAGTGGAGTTA | NR_029655.1 | Mir34b |
| **MIR34C** | AGTTACTAGGCAGTGTAGTTAGC | TTACCTGGCTGTGTGGTTAG | NR_029654.1 | Mir34c |
| **Mmp10** | ACGTACTTCTTTGTAGGGGACAA | TGTCTTGGGAAGCCTTTATCCA | NM_019471.2 | Mmp10 |
| **Mmp2** | CGAGGACTATGACCGGGATA | GGGCACCTTCTGAATTTCCA | NM_008610.2 | Mmp2 |
| **Mmp8** | ACGGTCTTCAGGCTGCTTA | AGCCACTTAGAGCCCAGTAC | NM_008611.4 | Mmp8 |
| **Mmp9** | TCCCCAAAGACCTGAAAACC | GGGTGTAACCATAGCGGTAC | NM_013599.2 | Mmp9 |
| **Nfkb1** | ACCGTATGAGCCTGTGTTCA | GTAGCCTCGTGTCTTCTGTCA | NM_008689.2 | Nfkb1 |
| **Nog** | AGCAAGAAGCTGAGGAGGAA | TAGGTCATTCCACGCGTACA | NM_008711.2 | Nog |
| **Pdgfa** | TGTAACACCAGCAGCGTCAA | GGCTTCTTCCTGACATACTCCA | NM_008808.3 | Pdgfa |
| **Runx2** | TCTGGCCTTCCTCTCTCAGTAA | AACTGCCTGGGGTCTGAAAA | NM_009820.4 | Runx2 |
| **SATB2** | CCAGGAGTTTGGGAGATGGTA | TGAAAGGTTCTCTCGCTCCA | NM_139146.2 | Satb2 |
| **SF3B4** | ATGCCCAAGGACAGAGTCAC | TGGCATAGTCGGCATCTTCC | NM_153053.4 | Sf3b4 |
| **Smad1** | CTCAGCCCATGGACACGAA | CTCGTAAGCAACTGCCTGAAC | NM_008539.3 | Smad1 |

| | | | | |
|---|---|---|---|---|
| **Smad2** | ACAAGTGACCAACAGTTGAACC | CAGGAGAGAGAGTAGTAGGAGACA | NM_001252481.1 | Smad2 |
| **Smad3** | TGCAGCCGTGGAACTTACAA | AAAGACCTCCCCTCCGATGTA | NM_016769.4 | Smad3 |
| **Smad4** | CCAACATTCCTGTGGCTTCC | GCTATCTGCAACAGTCCTTCAC | NM_008540.2 | Smad4 |
| **Smad5** | CCCAGCCTATGGATACAAGCA | CTCATAGGCGACAGGCTGAA | NM_008541.3 | Smad5 |
| **Sost** | CTCCCCACCATCCCTATGAC | CTGTCAGGAAGCGGGTGTA | NM_024449.5 | Sost |
| **Sox9** | AGTACCCGCATCTGCACAA | GTCTCTTCTCGCTCTCGTTCA | NM_011448.4 | Sox9 |
| **Sp7** | AGGATGGCGTCCTCTCTG | AGAGCCGCCAAATTTGCT | NM_130458.3 | Sp7 |
| **SPP1** | TGCCTGACCCATCTCAGAA | AAGTCATCCTTTTCTTCAGAGGAC | NM_009263.3 | Spp1 |
| **Tgfb1** | GCTGCGCTTGCAGAGATTAA | GTAACGCCAGGAATTGTTGCTA | NM_011577.1 | Tgfb1 |
| **Tgfb2** | GCCCATATCTATGGAGTTCAGACA | AGCGGAAGCTTCGGGATTTA | NM_009367.3 | Tgfb2 |
| **Tgfb3** | TCAGGCCCTTGCCCATAC | CTCTGGGTTCAGGGTGTTGTA | NM_009368.3 | Tgfb3 |
| **Tgfbr1** | AATTGCTCGACGCTGTTCTA | ACCGATGGATCAGAAGGTACA | NM_009370.2 | Tgfbr1 |
| **Tgfbr2** | TCTGTGAGAAGCCGCATGAA | GGCAAACCGTCTCCAGAGTAA | NM_009371.3 | Tgfbr2 |
| **Tgfbr3** | TGGTGTGGCATGTGAAGACA | GATGAAAACTGGACCACAGAACC | NM_011578.3 | Tgfbr3 |
| **Trp53** | CACAGCGTGGTGGTACCTTA | CCCATGCAGGAGCTATTACACA | NM_011640.3 | Trp53 |
| **Twist1** | CATGTCCGCGTCCCACTA | TGTCCATTTTCTCCTTCTCTGGAA | NM_011658.2 | Twist1 |
| **Vegfa** | CCAGCACATAGGAGAGATGAG | CTGGCTTTGTTCTGTCTTTCTT | NM_001025250.3 | Vegfa |
| **Vegfb** | GCCCCAGCCACCAGAA | GCTGGGCACTAGTTGTTTGAC | NM_001185164.N | Vegfb |
